# Robust Deep Denoising of Soft X-Ray Tomography Data for Biological Research: Targeting Tilt Series Versus Reconstructed Tomograms

**DOI:** 10.1101/2025.05.16.654457

**Authors:** ShaoSen Chueh, Jeremy C. Simpson, Sergey Kapishnikov

## Abstract

1.

Soft X-ray microscopy (SXM) is a powerful tool for nanoscale 3D imaging of hydrated, intact cells. However, its application is limited by pixel-wise-correlated noise inherent in the imaging process. While the deep-learning-based framework Noise2Inverse has been proposed for tomography denoising, the lack of robust validation for newly revealed features raises concerns regarding their practical biological utility. We argue that denoising the raw tilt series, instead of on the reconstructed tomogram, offers a significant advantage by leveraging the tomographic reconstruction process to mitigate local prediction errors through averaging. Consequently, features present in the tomogram reconstructed from a denoised tilt series exhibit higher credibility, as falsely predicted details on certain tilt series slices are likely to be averaged out. Nevertheless, denoising the tilt series remains challenging due to the spatially correlated noise that occurs in bioimaging. Existing self-supervised methods relying on a single noisy image struggle with such noise, while the Noise2Noise framework necessitates paired noisy datasets for training. This study addresses these challenges by investigating practical imaging workflows in SXM and related bioimaging modalities to identify existing imaging resources as training data for tilt series denoising. We compare the denoising performance when applied to tilt series versus reconstructed tomograms and evaluate the efficacy of chosen methods on real biological specimens to assess their applicability within practical SXM research. Our findings establish a robust and efficient denoising strategy for SXM by directly addressing the complexities of noise in the raw tilt series.

**Highlights:**

- Targeting tilt series for denoising inherently validates the correctness of the denoised tomogram via the reconstruction process.
- Existing multi-frame imaging schemes provide necessary resources for training Noise2Noise framework for tilt series denoising.
- Noise2Noise applied to tilt series reveals finer and more reliable details than direct denoising on reconstructed tomograms (Noise2Inverse).
- Denoising tilt series minimizes inference time compared to Noise2Inverse in soft X-ray tomography.

## 2. Introduction

Noise generated during the imaging process is a common problem across various imaging modalities. Many attempts have been made to tackle the denoising problem. Spatial filtering methods [1] are the most common methods with easy implementation. However, filtering-based methods tend to introduce blurring effects that could affect important information within the image. For example, BM3D [2] uses block-matching and 3D filtering to achieve denoising. However, it introduces certain artefacts during denoising, and also loses fine details. With the rapid development of deep learning, many neural network-based approaches are being explored. However, in practical bioimaging imaging settings, clean images are seldom available for training supervised deep learning models. Noise2Noise [3] tries to use a pair of noisy images with the same true signal to achieve similar performance as those with clean images. However, several assumptions regarding the noise distribution are still to be clarified more explicitly in the later section. Afterwards, more self-supervised methods were proposed that only require a single noisy image. For instance, Noise2Void (N2V) [4] enables denoising from a single noisy input but relies on the strict assumption of pixel-wise independent noise. The authors noted a performance degradation of N2V when applied to data exhibiting dependent noise. Recorrupted-to-Recorrupted (R2R) [5] (Pang et al., 2021), on the other hand, is a self-supervised model that recorrupts a single noisy image with 2 independent noise, and follows the N2N principles for model training. However, it is highly dependent on the accuracy of the assumed noise model, and any mismatch between the simulated and actual noise distribution can lead to suboptimal denoising performance or artefacts in the output, as shown in our experiment results. Other self-supervised methodologies, such as Neighbor2Neighbor [6], Self2Self [7], and Noise2Self [8] have also been proposed, each underpinned by specific assumptions that may not be valid for the noise characteristics inherent in soft X-ray microscopy images.

The Noise2Inverse framework [9] addresses self-supervised tomography denoising by splitting the noisy sinograms into multiple subsets. Each of these subsets is then independently reconstructed, resulting in multiple sub-tomograms that contain the same underlying clean signal but with different realizations of the noise. These sub-tomograms, depicting the same object, are then used to train a denoising model following the Noise2Noise principles. However, the major disadvantage of directly denoising the reconstructed tomogram is that there is a lack of mechanism to validate the correctness of the denoised output, which makes the model less credible for applications to real biological research. Moreover, the splitting operations will likely lose some resolution in the reconstructed sub-tomograms. Although the Noise2Inverse work demonstrates effective denoising tomograms of medical CT and foam datasets, SXM is more resolution-sensitive that its resolution range is on the edge of resolving small organelles. Lastly, the inference time could be long due to the large number of slices in reconstructed SXM tomograms (1024 or 2048 slices) without a decent GPU, which is usually the case for a biology laboratory.

In this research, different from Noise2Inverse, we focus on denoising the tilt series. The most important advantage of denoising the tilt series instead of the reconstructed tomogram is that it can better ensure the correctness of the noise prediction. If the model makes an error prediction on a certain tilt series slice, it is unlikely the same error will happen across all tilt series slices. Thus, the single mistake can be averaged out during the reconstruction by the signals on other slices instead of making up novel details on the reconstructed tomograms. Furthermore, targeting tilt-series can minimize the inference time compared to the reconstructed tomograms. The number of tilt series slices is usually 111-141 or fewer, depending on the accessible tilt angle, while the number of slices in a SXM tomogram is usually 1024 or larger. Thus, we argue that tilt-series is a better denoising target for SXM compared to reconstructed tomograms as used in Noise2Inverse.

To enable the training of denoising models on the tilt series, we investigate current SXM imaging practices and identify that multiple frames at a single tilt angle are usually collected. This multi-frame collection scheme is also common in other bioimaging modalities such as cryo-electron microscopy (cryo-EM) [10] and movie-mode dynamic electron microscopy [11]. Thus, this work highlights the substantial potential of leveraging existing imaging data to support deep learning-based denoising approaches. Our objective is to assess the practicality of denoising the tilt series and to compare its implementation efficiency and denoising performance with that of tomogram-based methods, such as Noise2Inverse. Finally, we validate our findings on real biological datasets to evaluate their effectiveness and applicability in realistic SXM imaging workflows.

## 3. Data

### 3-1. Tilt Series Acquisition and Reconstruction

The 3D soft X-ray tomography is reconstructed with a series of 2D projections called “tilt series”. The tilt series is acquired by using a soft X-ray light source to image biological samples with angles approximately +/-50 – 70 degrees (Figure 1). After aligning the tilt series, these 2D projection slices can be used to reconstruct a 3D tomography of the biological sample (Figure 2).

**Figure 1.**
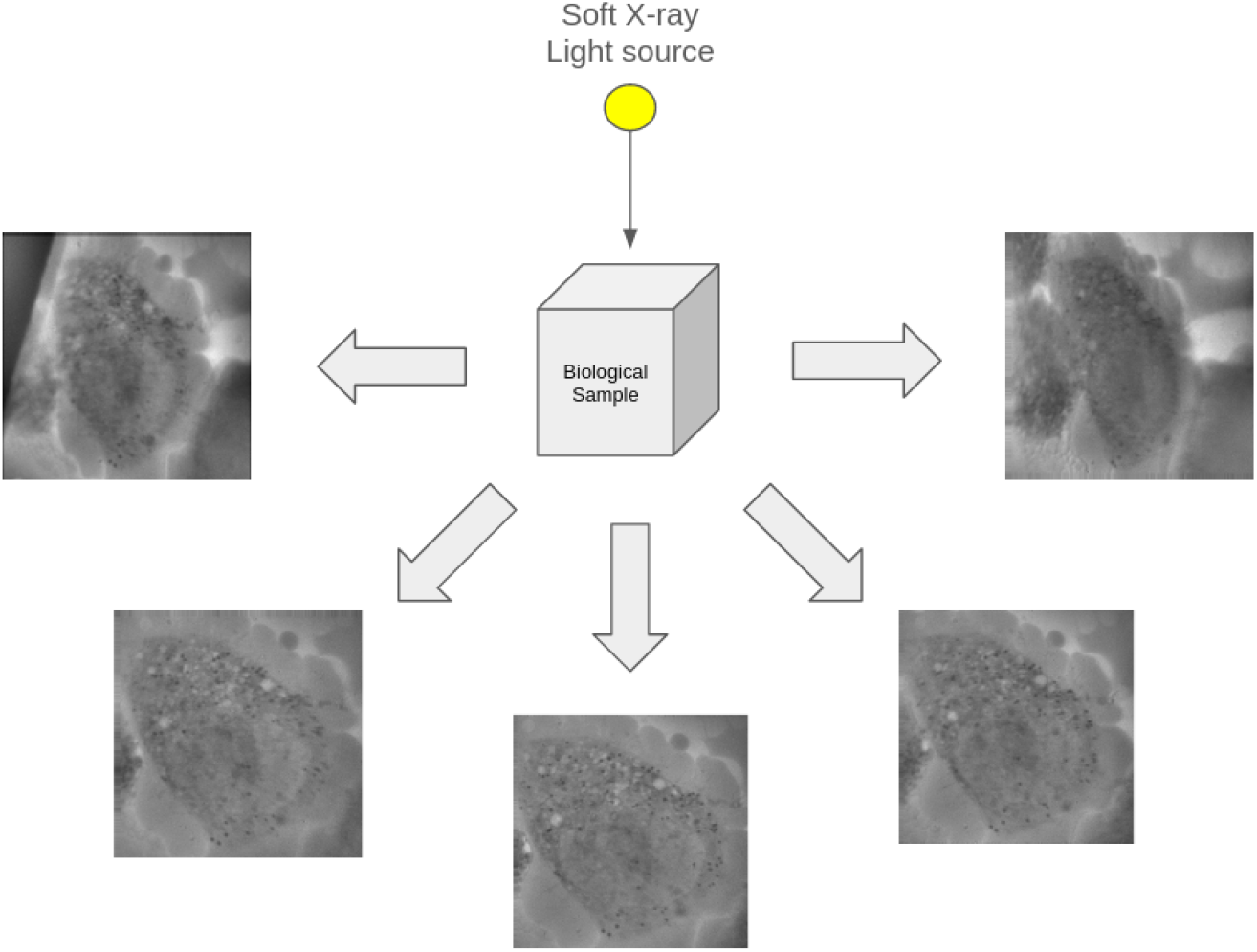
2D Tilt Series Acquisition. A soft X-ray light source is passed through the specimen to acquire a stack of 2D projections (tilt series) with an approximate angle of +/-50 – 70 degrees.

**Figure 2.**
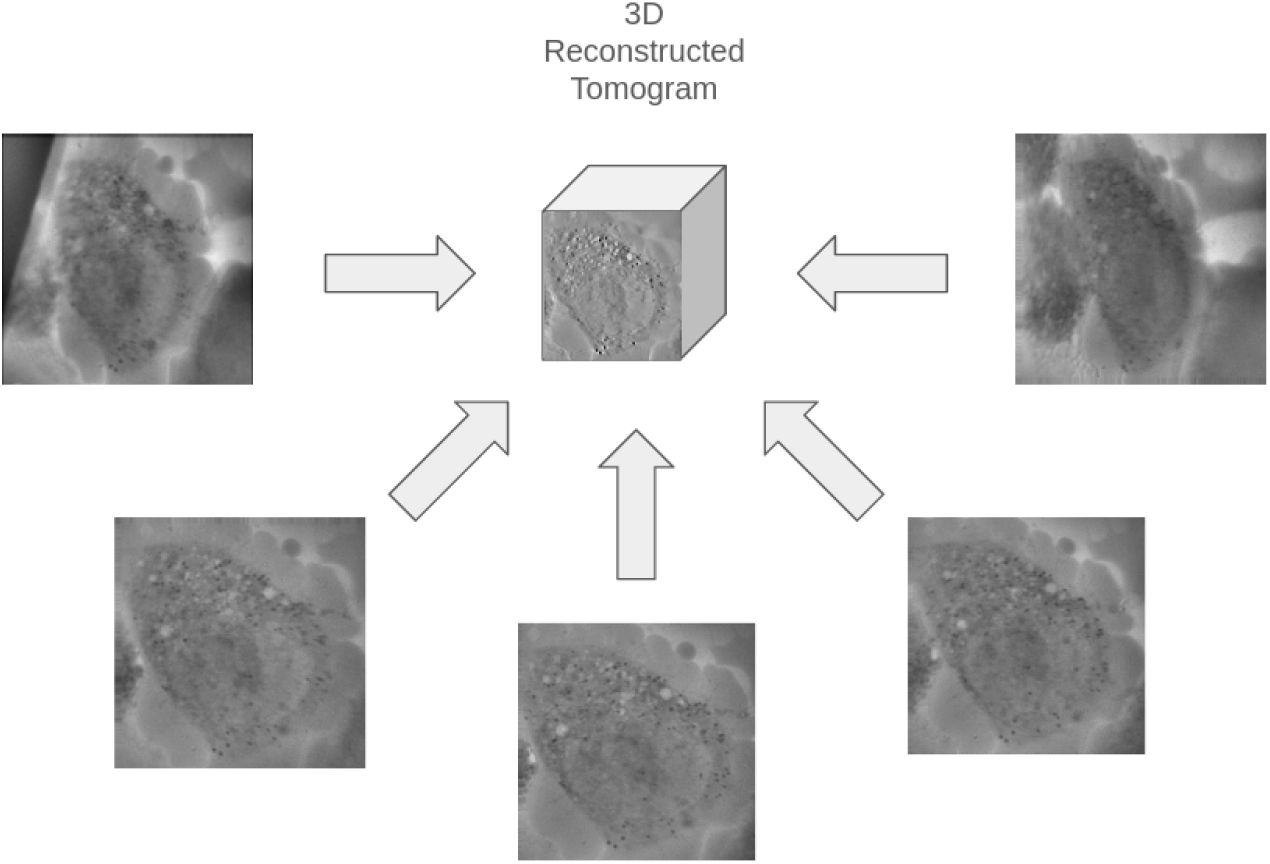
3D Tomography Reconstruction. The acquired tilt series is used to reconstruct a 3D tomogram of the biological sample. Common reconstruction algorithms include Weighted Back Projection (WBP), Simultaneous Iterative Reconstruction Technique (SIRT), and Deep-Learning-Based algorithms.

### 3-2. Multi-Frame Imaging and Shading Correction

In SXM, tilt series projections from a single degree can be acquired in multiple frames per tilt, particularly for lab-based microscopes in order to minimize the contribution of the thermal drifts, where the exposures could be long. In every tilt, the frames are subsequently registered to each other and summed up to a single frame. This multiframe data collection scheme provides the required input for the Noise2Noise framework as these duplicates acquired from the same degree contain the same sample with pairwise independent noise.

Nonetheless, during acquisition, the soft X-ray illumination experiences drifts (Figure 3). The illumination drift (part of the true signal) in the images needs to be corrected to ensure the consistency of the true signal in the duplicate frames.

**Figure 3.**
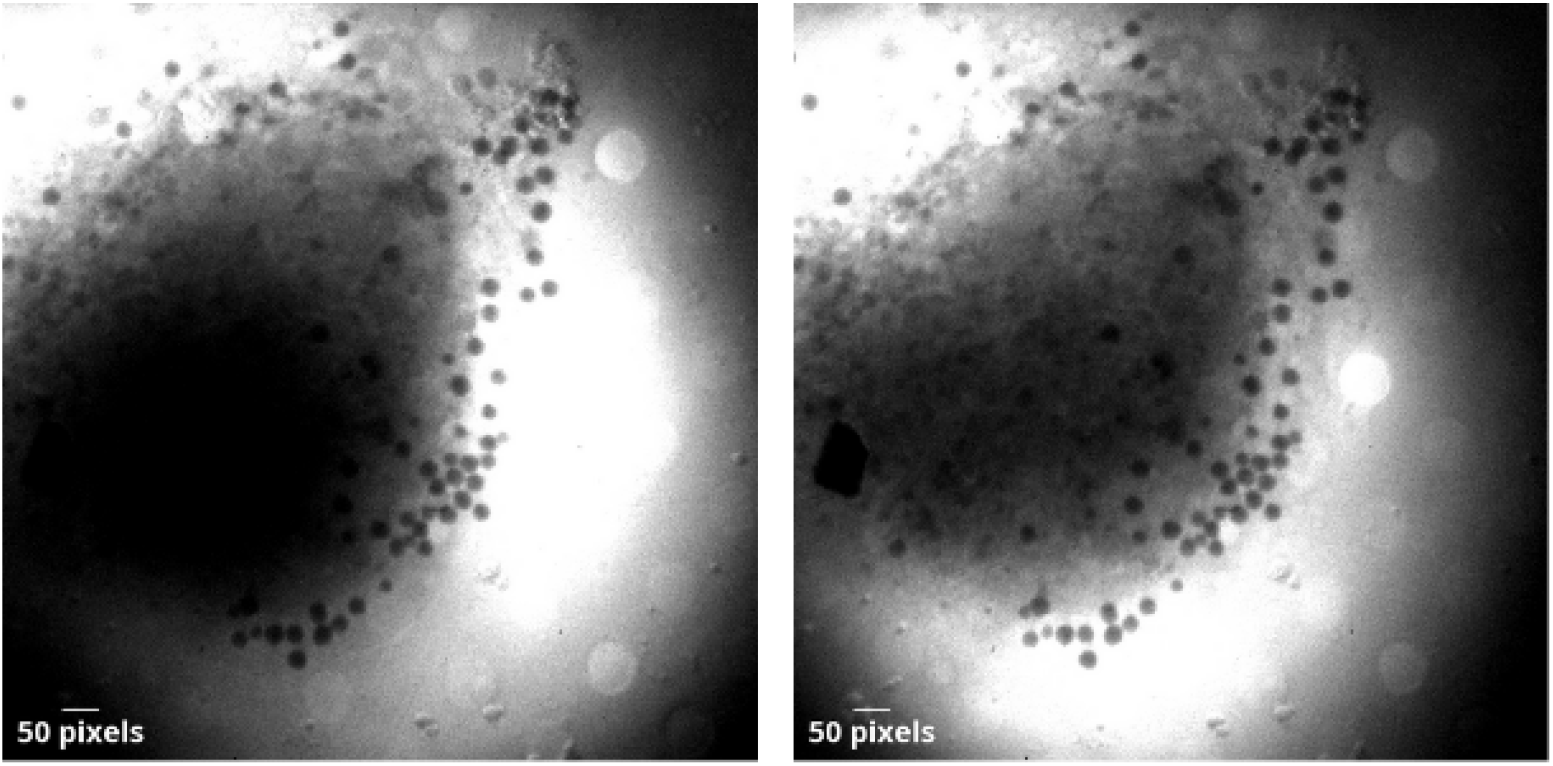
Multi-frame imaging with illumination drift **(Pixel size: 28.8 nm)**. Due to the inevitable illumination drift, the duplicate frames acquired from the same perspective still demonstrate differences in the underlying signal, which damages the signal assumption of the Noise2Noise (N2N) framework.

However, except the illumination difference, the high-frequency signals of the duplicate frames are still consistent. Therefore, we propose using the gaussian high-pass filtering to correct the differences in illumination (Figure 4).

**Figure 4.**
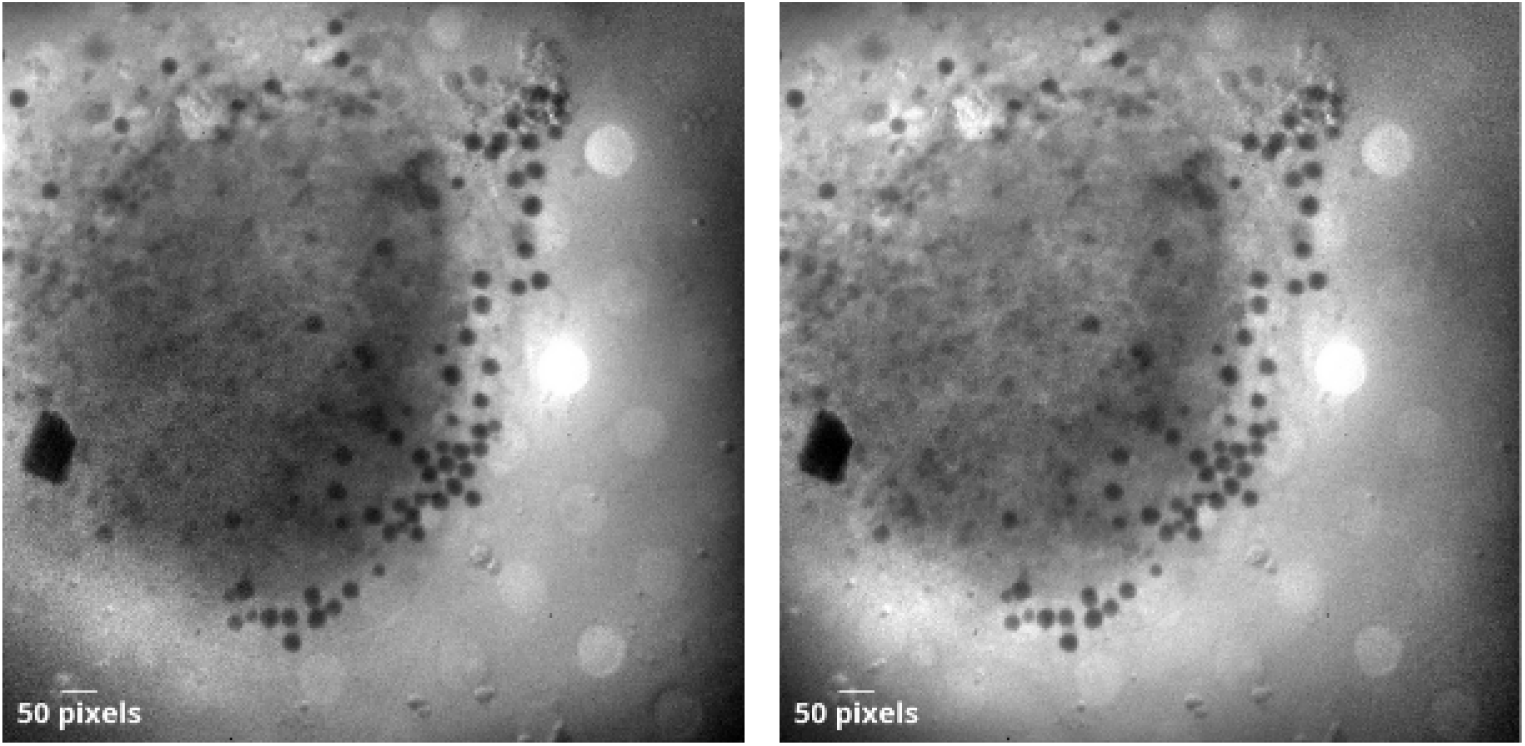
Gaussian high-pass filtering **(Pixel size: 28.8 nm)**. The high-pass filter effectively removes the difference in low-frequency signals (illumination), and thus allows the high-frequency features to be consistent across duplicate frames.

The major problem of this shading correction method is that the filtered images lose a certain level of contrast, and this causes some difference in low-frequency signal between the training data (filtered duplicate frames) and original real data, which could potentially cause issues for deep learning models to generalize on real images. However, in our experiments, we found the model still achieves a satisfactory denoising performance in inference time even with the differences in low-frequency signals.

### 3-3. Noise Analysis

The noise in soft X-ray microscopy arises from three primary sources: (1) Poisson noise due to the stochastic arrival of X-ray photons at the detector, (2) scattered radiation noise from unwanted non-image forming photons that reduce the overall contrast, and (3) structural noise introduced by the detector itself, exhibiting spatial correlation patterns. However, because we are unable to isolate the noise from the signal due to the fact that the clean image does not exist, definitively quantifying the specific characteristics of the noise requires further investigation or alternative methodologies.

On the other hand, we can qualitatively observe certain noise patterns in the soft X-ray images to confirm that the noise is pixel-wise-correlated (Figure 5).

**Figure 5.**
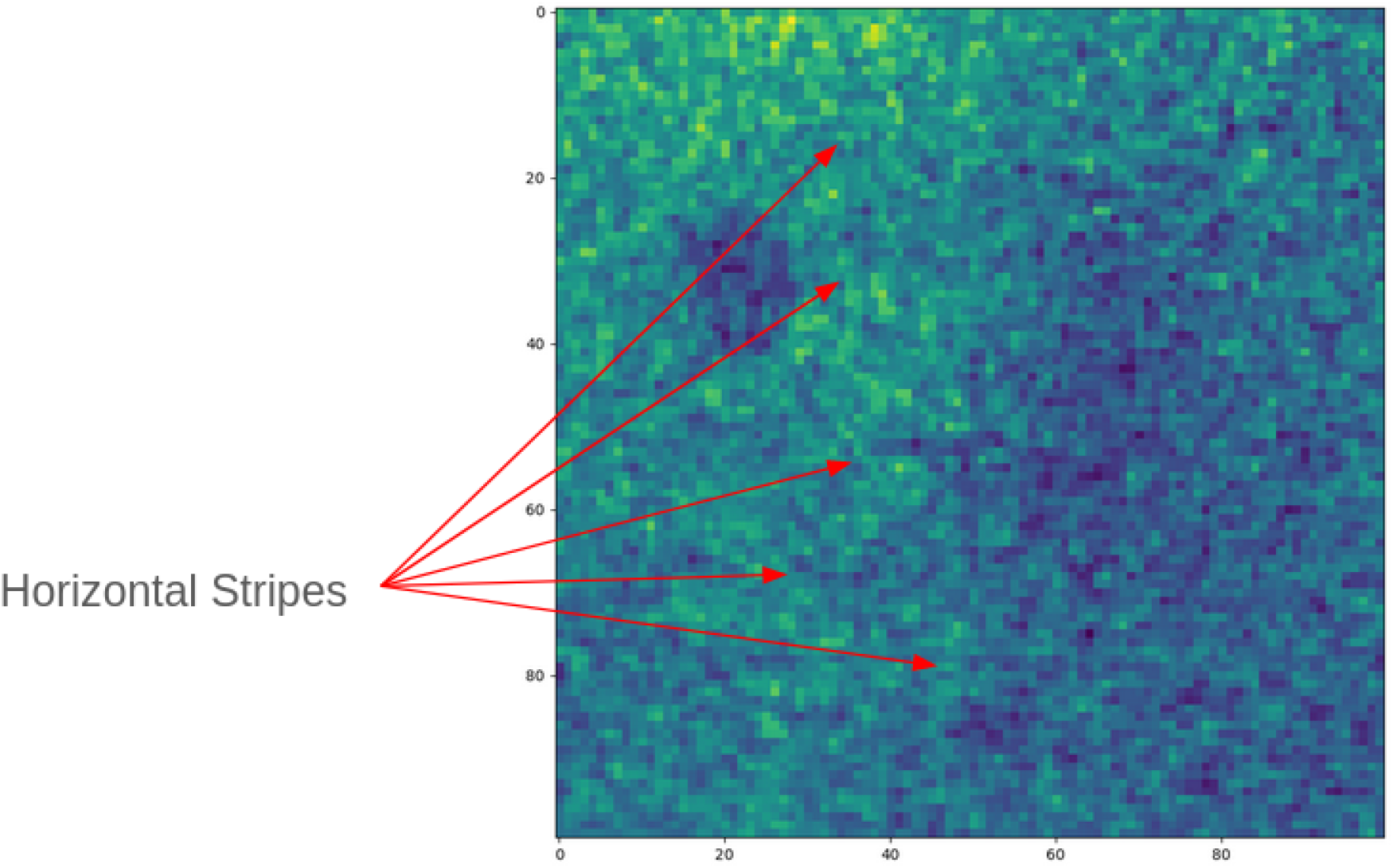
Spatially-correlated noise patterns. The spatially-correlated noise (horizontal and vertical stripes) are likely introduced by the detector.

### 3-4. Sample Source and Imaging

In this study, two types of samples were used for demonstration: Human Pulmonary Fibroblast (HPF cells and A549 cells. Both cell types were grown on 3 mm TEM grids (Au G200F1 with Quantifoil R2/2 film) and vitrified by plunge-freezing using either a Leica GP2 or an FEI Vitrobot Mark IV plunge-freezer. Sample preparation was carried out at the Center for Optical Technologies, Aalen University, and the Core Facility for Integrated Microscopy (CFIM) at the University of Copenhagen. The vitrified samples were subsequently imaged using soft X-ray cryo-tomography in the laboratory-based soft X-ray microscope SXT-100, developed by SiriusXT.

Tomograms were acquired by projecting soft X-rays through the samples over tilt series typically ranging from -60° to +60° in 1° increments. Each projection was recorded using a multi-frame acquisition scheme (2 - 3 frames per tilt, 15 - 30 seconds exposure per frame). Frames at each tilt angle were aligned to one another (e.g., using cross-correlation) to remove potential thermal drift effects.

The data referenced in this study are available from the corresponding author on request.

## 4. Methodology

### 4-1. Noise2Noise Framework

In the ideal setting, a pair of noisy (*x*) and clean (*y*) images are required to train a supervised denoising model as the expression :

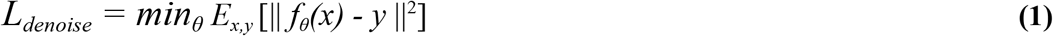

for *f*_*θ*_() represents a parametric neural network. However, in many situations the clean image targets do not exist, including in this research. On the other hand, Noise2Noise argues that the clean target image is not necessary. Instead, a noisy pair of images (*x*_*1*_,*x*_*2*_) with the same underlying true signal (*y*) is sufficient. The objective for training of the Noise2Noise can be reformulated as

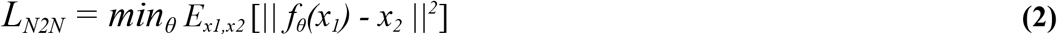

The arguments are that a noisy image (*x*) can be seen as the true signal (*y*) + the noise (*n*), and thus the objective can be expanded as

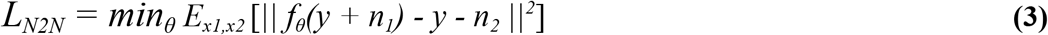

Assuming the noise(*n*) has zero mean, the objective expression is identical to the original supervised objective term :

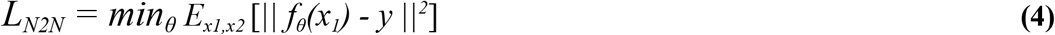

However, in the SXM tilt series, the statistics of the noise is complicated. It is a combination of Poisson noise and spatially-correlated (horizontal and vertical stripes) noise, and thus unlikely to meet the zero-mean assumption. Nonetheless, we could expand **(3)** into

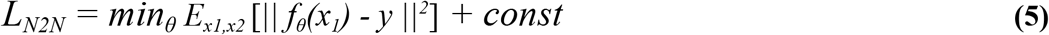

considering the non-zero-mean noise, which indicates that the model does not actually learn the true clean signal, but with a constant combining mean of the signal + the noise mean. However, we argue that this objective loss term can still provide satisfactory denoising performance, and the effect of the constant can be minimized with post-processings, as shown in our experiment results.

Another similar but more extreme case, text-removal in the N2N work, shows its capability of effective removal of spatially-correlated non-zero-mean noise. In the text removal task, there are 4 scenarios at each pixel of the paired noisy images:

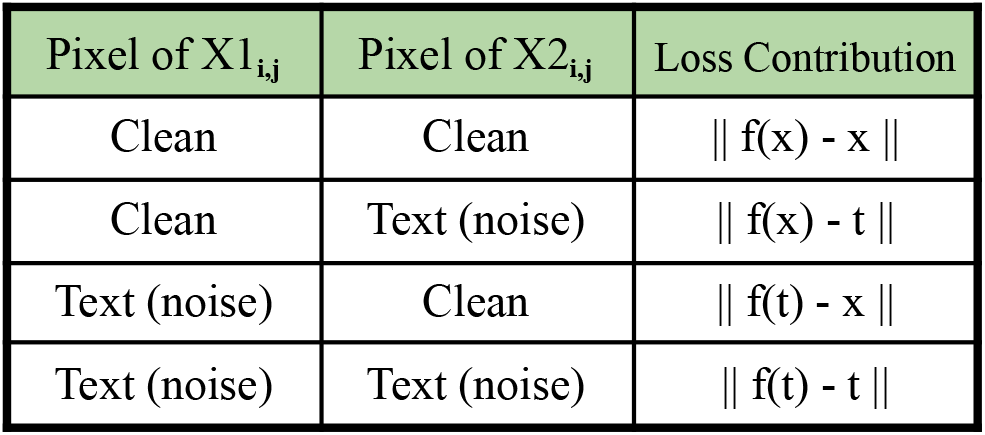

In some scenarios (text-text, clean-text), the loss objective seems to force the model to learn the undesired noise, but it is worth noting that the probability of a clean pixel is much higher than a text-corrupted pixel in the text removal case. As long as the structural noise (text) is added pairwise independently, the objective loss term (2) still motivates the model to approximate the underlying signal. Therefore, even if the noise is structural, non-zero-mean, and not pixel wise independent, as long as the noise does not overwhelm the true signal, nor pairwise correlated, the Noise2Noise framework can still provide reasonable denoising performance.

### 4-2. Model Architecture

In this research we adopt Denoising Convolutional Neural Network (DnCNN) [12] as the backbone for the Noise2Noise framework. The model is designed to predict a noise component from a noisy input, which is subsequently subtracted to obtain the denoised image. The architecture consists of an initial convolutional layer with 128 feature maps followed by a ReLU activation. This is followed by 15 intermediate layers, each comprising a convolutional layer (kernel size 3×3, padding 1), batch normalization, and ReLU activation. A final convolutional layer maps the output back to the input channel dimension. In total, the network contains 17 layers. The output is optionally normalized to the range [0, 1] to improve training stability and facilitate qualitative evaluation. All convolutional layers use a kernel size of 3×3 with zero-padding to preserve spatial resolution.

## 5. Results

For evaluation, we implement both Noise2Noise (N2N, on tilt series) and Noise2Inverse (N2I, on reconstructed tomogram) methods to compare their performance. Furthermore, a series of self-supervised frameworks are also implemented (N2V, NBR2NBR, R2R, on tilt series) to explore the feasibility of single-slice denoising of spatially-correlated noise on tilt series. Our results contain both qualitative and quantitative evaluation. However, it should be noted that because clean target images do not exist, we are unable to separate the noise and the target, and thus the quantitative analysis can only be used as reference for perceived quality of the images. Therefore, greater emphasis is placed on qualitative analysis when observing the performance on real biological sample images.

### 5-1. Qualitative Analysis

Qualitative analysis reveals effective denoising by N2N, N2I, contrasting with the suboptimal performance of self-supervised methods (NBR2NBR, R2R, N2V) likely due to violated noise distribution assumptions inherent in SXM images (Figure 6, Figure 7). Both N2V and NBR2NBR assume independent noise from the original image. However, as shown in the noise analysis section, SXM image noise is spatially correlated, and thus causes the degradation of the performance. On the other hand, R2R does not assume independent noise from the original image. However, since we are unable to accurately quantify and simulate the SXM noise, simply adding Gaussian noise to recorrupt the original image cannot generalize to real SXM images, and thus fails to perform as good as other methods.

**Figure 6.**
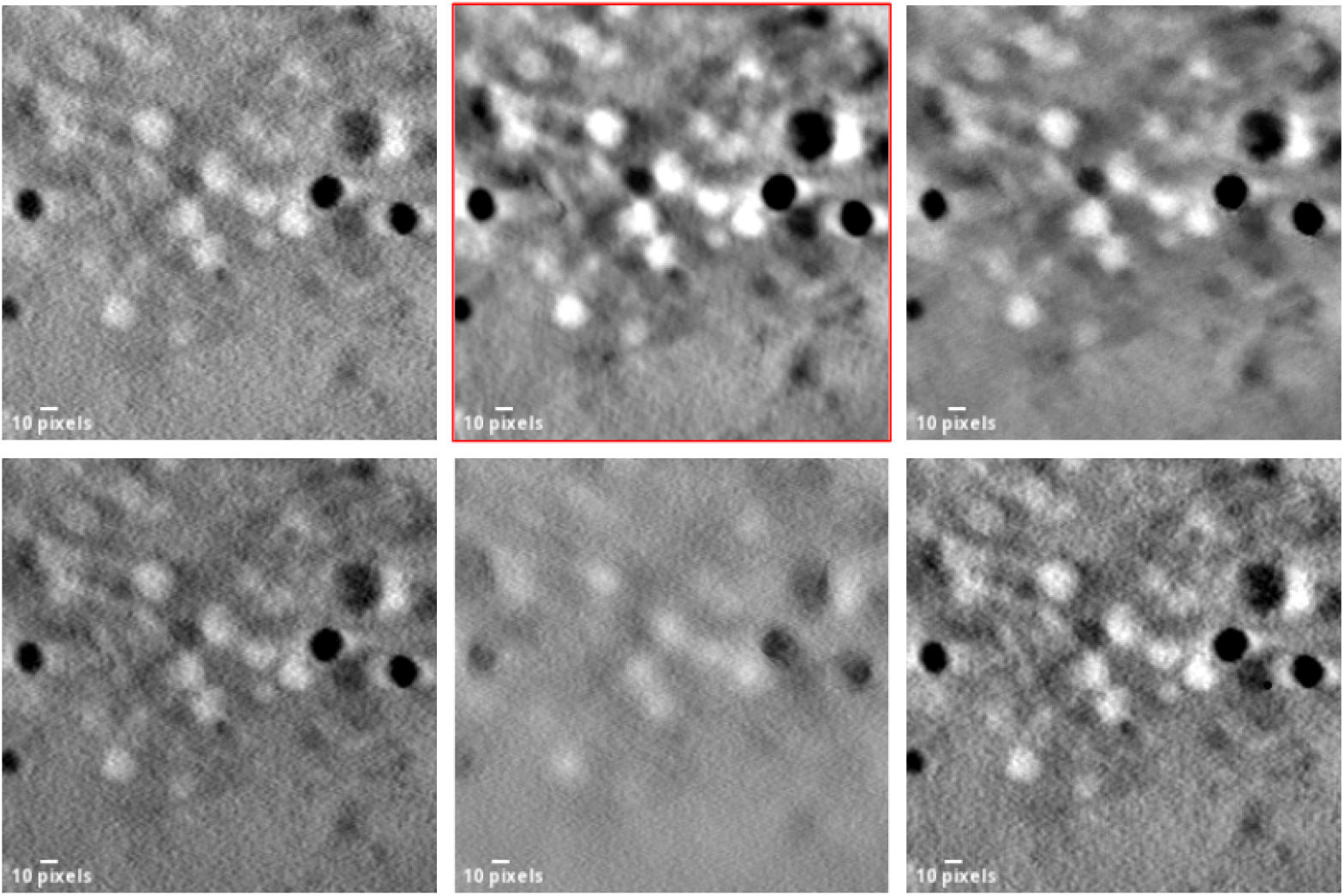
Denoised tomogram slice. **ROW1** : original, N2N, N2I. **ROW2** : NBR2NBR, R2R, N2V. **(pixel size : 25.3 nm)**. The denoised output indicates the superior denoising performance of N2N and N2I compared to self-supervised frameworks, due to the unmet pixel-independent noise assumption. Further comparison between N2N and N2I is shown in Figure 8.

**Figure 7.**
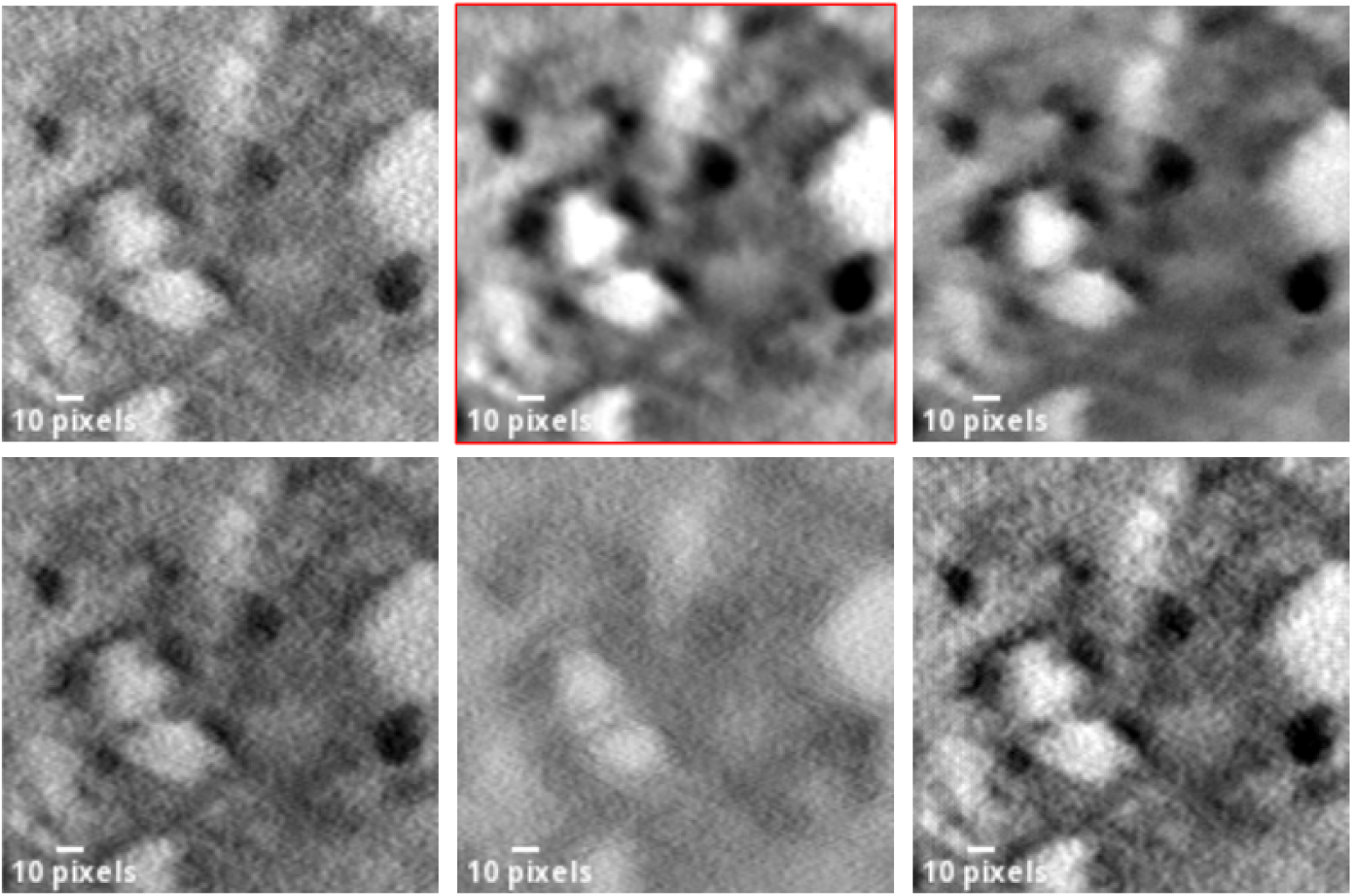
Denoised tomogram slice. **ROW1** : original, N2N, N2I. **ROW2** : NBR2NBR, R2R, N2V. **(pixel size : 25.3 nm)**. The denoised output indicates the superior denoising performance of N2N and N2I compared to self-supervised frameworks, due to the unmet pixel-independent noise assumption. Further comparison between N2N and N2I is shown in Figure 8.

While N2I denoising performance is comparable to N2N, it is observed that N2I tends to smooth out certain details, resulting in some blurry effects, compared to N2N (Figure 8). This situation could potentially result from the sinogram-splitting operations.

**Figure 8.**
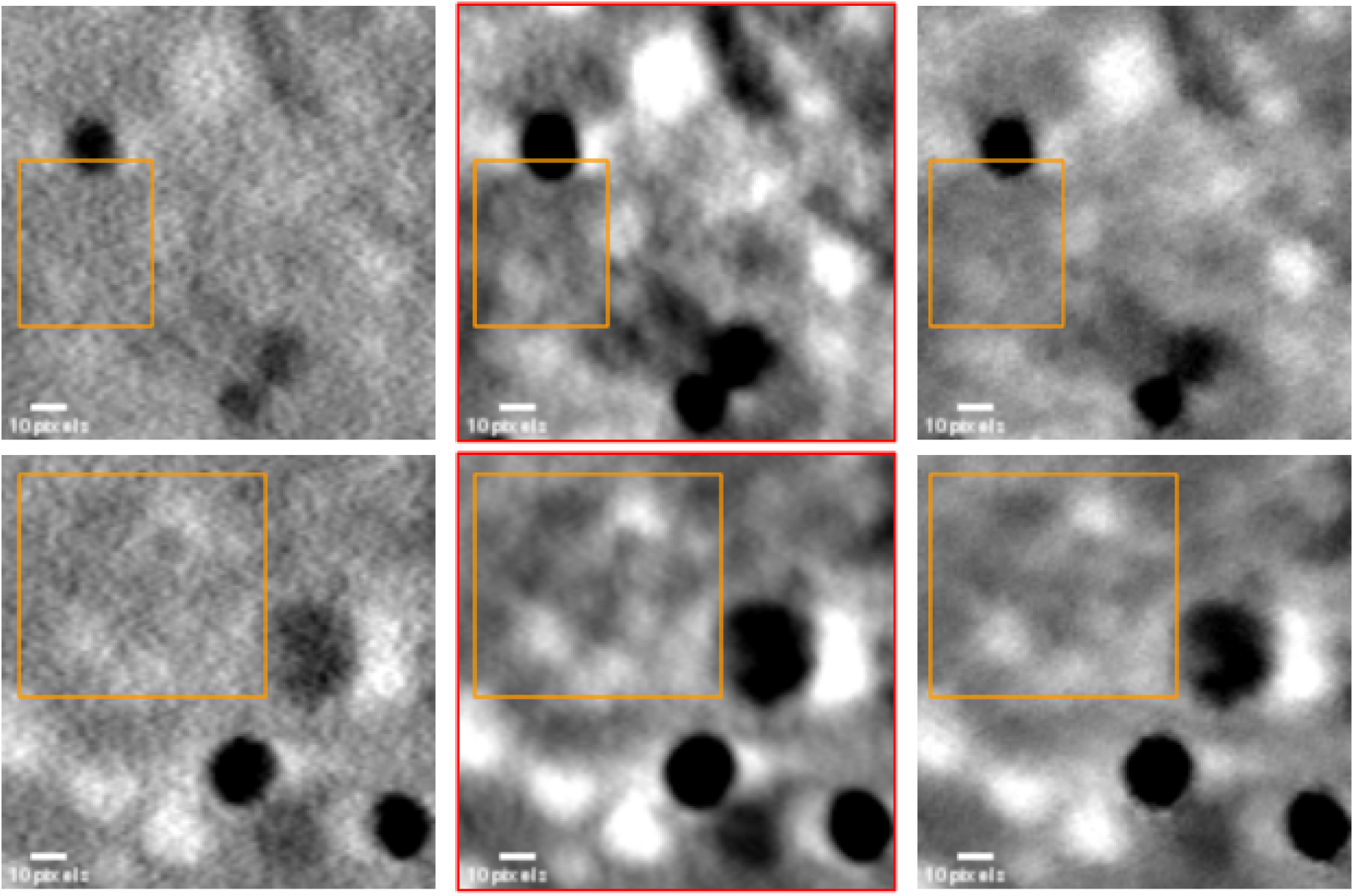
Original(Left Column), **N2N**(Mid Column), N2I (Right Column). **(pixel size : 25.3 nm)**. The N2N framework targeting the tilt series reveals reliable and clearer features (orange boxes), while N2I targeting the reconstructed tomogram tends to over smooth details. This over-smoothing effect of N2I could result from the sinogram splitting mechanism for generating paired sub-tomograms for training, which leads to the loss of a certain degree of resolution.

N2I relies on the reconstructed sub-tomograms for training the denoising model. However, sub-tomograms reconstructed from sub-stacks of tilt-series lose a certain degree of resolution. Thus, the N2I model trained on lower-resolution data could potentially have difficulty recovering the fine details in inference time.

Most importantly, denoising the tilt series enhances the robustness of the output reconstructed tomograms. The purpose of performing denoising tasks is to reveal more information from the image. However, when the model does reveal further subtle details (Figure 8, 2nd row, orange boxes) that were unseen from the original image, it can be confusing for practical biological research. Therefore, it is important to have a validation mechanism to verify the existence of these features, and the reconstruction procedure of tomogram from tilt series serves as a great validation mechanism. This is because if the model falsely predicts the noise in a single tilt image, it is unlikely that the same error will consistently occur across all slices in the tilt series. Consequently, such isolated errors are averaged out during the reconstruction process, typically resulting in local blurring effect rather than the introduction of false structures. Therefore, when specific features appear in the reconstructed tomograms, even if somewhat inevident, their presence is highly indicative of true underlying structures. In contrast, direct denoising of reconstructed tomograms, as implemented in Noise2Inverse (N2I), lacks this intrinsic verification mechanism. Thus, we argue that this robustness to error propagation represents a key advantage of performing denoising on the tilt series.

Lastly, denoising tilt series could also potentially reduce imaging time. In SXM imaging, multiple frames of projections per tilt are usually averaged to acquire a better quality slice. In our experiments, we denoise a single frame projection (Figure 9, right) to compare it with the original single-frame projection (Figure 9, left) and the projection averaged with 2 frames (Figure 9, middle).

**Figure 9.**
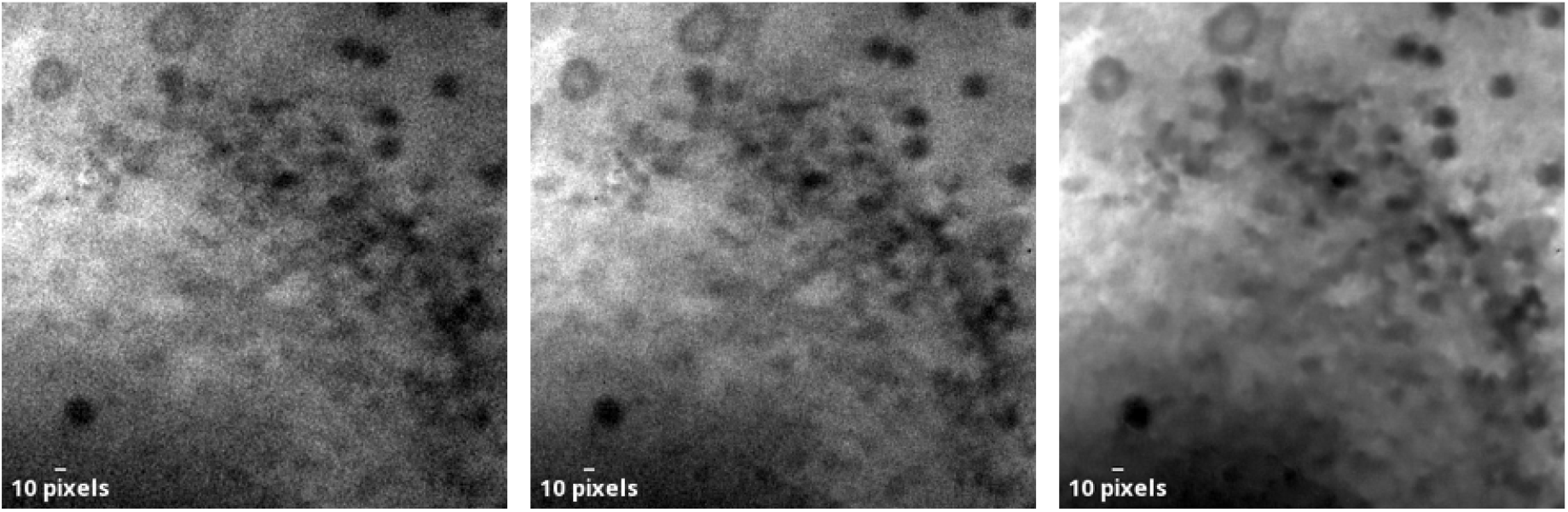
Original single projection (left). Averaged projection : 2 slices (mid). Denoised single projection (right). **(Pixel size: 28.8 nm)**. Even with multi we-frame averaging, the averaged slice still contains a certain level of noise. By contrast, the N2N framework effectively removes the noise with only a single slice.

It can be observed that the projection denoised from a single frame is of better quality than the average of 2 frames. This means that it could halve the imaging time by acquiring only a single frame per tilt. Moreover, minimizing the duration of soft X-ray exposure through this approach can mitigate the risk of damage to the biological specimen.

### 5-2. Quantitative Analysis

To provide an objective assessment of denoising performance, we employed two No-Reference Image Quality Assessment (NR-IQA) metrics: BRISQUE (Blind/Referenceless Image Spatial Quality Evaluator) [13] and PIQE (Perception-based Image Quality Evaluator) [14]. Additionally, we utilized Fourier Ring Correlation (FRC) to estimate the resolution of the reconstructed tomograms. The inherent absence of a clean ground truth in soft X-ray microscopy complicates a definitive quantitative evaluation. Furthermore, it is important to acknowledge that each metric possesses inherent biases that limit a completely unbiased performance assessment. Therefore, we implemented these three metrics to gain a more comprehensive understanding of the perceptual quality and resolution of the denoised images.

BRISQUE operates by extracting statistical features based on natural scene statistics, positing that noise and artefacts disrupt these inherent regularities. This method relies on a Support Vector Machine (SVM) trained on natural images, potentially introducing bias. PIQE, conversely, directly quantifies perceptual quality through an unsupervised analysis of local spatial intensity variations, identifying deviations in distortion and contrast from typical natural image characteristics. FRC estimates resolution by assessing the cross-correlation of Fourier transforms from two independently reconstructed sub-tomograms derived from split tilt series, providing a measure of reproducible high-frequency content.

Our NR-IQA results (Figure 10) indicate that Noise2Noise (N2N) applied to the tilt series outperforms Noise2Inverse (N2I) in terms of PIQE score, suggesting better perceptual quality according to this metric. Conversely, N2I achieved a superior BRISQUE score. Notably, both N2N and N2I demonstrated a considerable performance margin over other self-supervised denoising methods, implying a higher perceptual quality of their outputs compared to these methods and the original tomogram.

**Figure 10.**
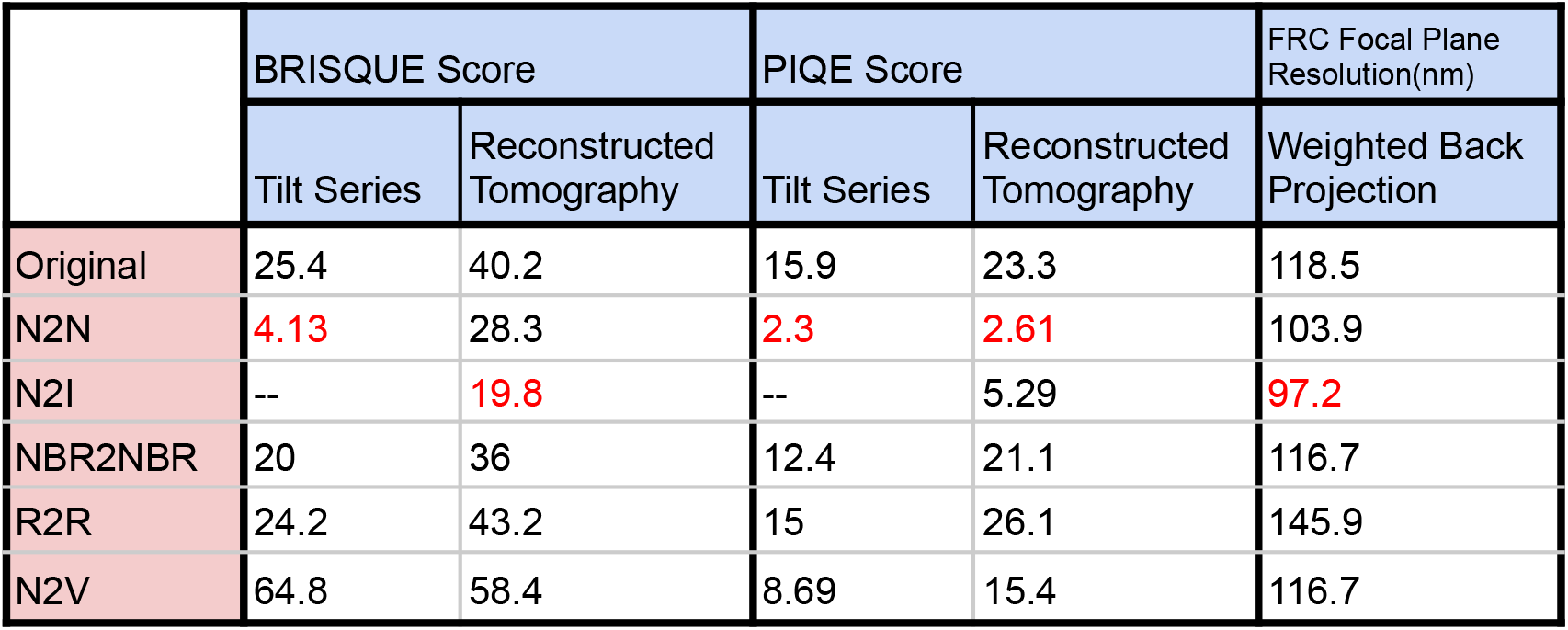
Evaluation metrics (*red text indicates the best score achieved*). Both N2N and N2I demonstrate superior performance compared to other self-supervised frameworks in terms of image quality. It should be noted that each quantitative metric possesses inherent biases to varying extents. Consequently, these metrics should be considered as supplementary references rather than definitive determinants of overall evaluations.

The FRC analysis (Figure 10) revealed a substantial improvement in estimated resolution for both N2N and N2I compared to the other methods. Surprisingly, the FRC results indicated a slightly higher estimated confocal plane resolution for N2I than N2N, which contradicts our qualitative observations. We hypothesize that this discrepancy may arise from artefacts potentially introduced during the tomographic reconstruction of denoised tilt series, a process bypassed when directly denoising the reconstructed tomogram as in N2I.

In conclusion, the quantitative analysis, while constrained by the absence of a clean ground truth and inherent metric biases, suggests that both N2N applied to tilt series and N2I on reconstructed tomograms demonstrate significant improvements in perceptual quality and resolution. The subtle differences observed between the two methods, particularly in FRC, appear to be influenced by the distinct denoising procedures and the reconstruction algorithms employed. Therefore, while both approaches offer effective denoising for SXM tomography, the choice between them may depend on specific application requirements and computational resources.

## 6. Conclusion

This research explores a robust denoising method that can be applied to practical biological research. Denoising tilt series leverages the tomographic reconstruction procedure as a reliable validation mechanism for the denoised output. This mechanism ensures the robustness of deep-learning-based denoising approaches, which is a major advantage for practical biological applications. Furthermore, the reduction of inference time compared to Noise2Inverse is also considerable, especially in the setting of a biology laboratory without sufficient computing resources. While N2I also demonstrates promising denoising performance directly on reconstructed tomograms, it is challenging to verify the existence of newly revealed details after denoising. Furthermore, the degradation of resolution of N2I due to the sinogram-splitting operations for reconstructing sub-tomograms as training data is also an issue for practical applications.

In terms of implementations, N2I possesses an advantage over N2N denoising on tilt series that it can directly be implemented with 1 single noisy tomogram, even though it requires complex preprocessing steps. However, the multi-frame imaging scheme is common in certain bioimaging modalities, and we argue that it is worth exploring these readily available imaging resources for implementing N2N denoising on tilt series, with easy implementation that avoids complex preprocessing steps. Overall, while denoising on both targets provide promising denoising performance, each method has their own advantages, depending on the resources available of the bioimaging modalities.

For future research, our experiments revealed that self-supervised methods that require only a single noisy image heavily depend on the assumption of independent noise. Consequently, these methods yielded sub-optimal results when applied to spatially-correlated noise, such as that encountered in SXM imaging. This outcome indicates a significant area for future research in self-supervised denoising techniques specifically designed for spatially-correlated noise. Success in this domain holds the promise of substantially reducing the resource demands for denoising tilt series and avoids complex implementations in N2I.

## 7. Declaration of generative AI and AI-assisted technologies in the writing process

Statement: During the preparation of this work the author(s) used Gemini by Google only to improve the quality and readability of writing. After using this tool/service, the author(s) reviewed and edited the content as needed and take(s) full responsibility for the content of the published article.

## 8. Acknowledgements

The authors gratefully acknowledge Professor Heloisa Nunes Bordallo (Niels Bohr Institute, University of Copenhagen), Professor Fábio Rocha Formiga (Department of Immunology, Aggeu Magalhães Institute (IAM), Oswaldo Cruz Foundation (FIOCRUZ), Erasmus + International Credit Mobility Program (European Commission)), and Professor Andreas Walter (Aalen University, Center for Optical Technologies/Aalen School of Applied Photonics) for providing samples for soft X-ray tomograms used in this research. We also thank Professor Rasmus Hartmann-Petersen and Dr. Martin Grønbæk-Thygesen (UCPH-Department of Biology) for preparing the A549 cells. We also thank Tillmann Hanns Pape at the Core Facility for Integrated Microscopy (CFIM) at University of Copenhagen for assistance with sample preparation.

## 10. Data Availability Statement

The data referenced in this study are available from the corresponding author upon reasonable request.

## 11. Author Contributions

ShaoSen, Chueh : Methodology, Formal Analysis, Software, Writing - original draft Jeremy C. Simpson : Supervision, Writing - review and editing Sergey Kapishnikov : Methodology, Supervision, Validation, Writing - review and editing, Project Administration

